# Aiming off the target: studying repetitive DNA using target capture sequencing reads

**DOI:** 10.1101/2020.12.10.419515

**Authors:** Lucas Costa, André Marques, Chris Buddenhagen, William Wayt Thomas, Bruno Huettel, Veit Schubert, Steven Dodsworth, Andreas Houben, Gustavo Souza, Andrea Pedrosa-Harand

**Affiliations:** Laboratory of Plant Cytogenetics and Evolution, Department of Botany, Federal University of Pernambuco, Recife-PE, Brazil; Max Planck Institute for Plant Breeding Research, Cologne, Germany; AgResearch, Plant Functional Biology, Ruakura, New Zealand; New York Botanical Garden, Bronx, New York, United States of America; Max Planck Genome Centre Cologne, Max Planck Institute for Plant Breeding Research, Cologne, Germany; Leibniz Institute of Plant Genetics and Crop Plant Research (IPK) Gatersleben, Seeland, Germany; School of Life Sciences, University of Bedfordshire, Luton, UK

**Keywords:** Genome Skimming, holocentric, Reduced-representation sequencing, RepeatExplorer, *Rhynchospora*, satellite DNA, transposable elements

## Abstract

- With the advance of high-throughput sequencing (HTS), reduced-representation methods such as target capture sequencing (TCS) emerged as cost-efficient ways of gathering genomic information. As the off-target reads from such sequencing are expected to be similar to genome skims (GS), we assessed the quality of repeat characterization using this data.
- For this, repeat composition from TCS datasets of five *Rhynchospora* (Cyperaceae) species were compared with GS data from the same taxa.
- All the major repetitive DNA families were identified in TCS, including repeats that showed abundances as low as 0.01% in the GS data. Rank correlation between GS and TCS repeat abundances were moderately high (*r* = 0.58-0.85), increasing after filtering out the targeted loci from the raw TCS reads (*r* = 0.66-0.92). Repeat data obtained by TCS was also reliable to develop a cytogenetic probe and solve phylogenetic relationships of *Rhynchospora* species with high support.
- In light of our results, TCS data can be effectively used for cyto- and phylogenomic investigations of repetitive DNA. Given the growing availability of HTS reads, driven by global phylogenomic projects, our strategy represents a way to recycle genomic data and contribute to a better characterization of plant biodiversity.

## INTRODUCTION

One intriguing aspect of eukaryotic genomes is the staggering 60,000-fold variation in DNA content among species (Elliott & Gregory, 2015). Advances in genomics have led to the discovery that most of this diversity is the result of variable amounts of repetitive DNA, commonly divided into tandem repeats and dispersed repeats (Weiss-Schneeweiss *et al.*, 2015; Elliott & Gregory, 2015). Tandemly distributed satellite DNAs are remarkable for their fast evolution and variance in abundance and structure at all hierarchical levels (Novák *et al.*, 2017; Ávila Robledillo *et al.*, 2018). As for dispersed repetitive sequences, transposable elements (TE) are especially abundant in flowering plant genomes (Jurka *et al.*, 2011; Galindo-González *et al.*, 2017). Retrotransposons, particularly the ones possessing a long terminal repeat (LTR-retrotransposons) account for most of this abundance (Weiss-Schneeweiss *et al.*, 2015) with two major superfamilies being recognized (Ty1/copia and Ty3/gypsy) based on the order of the protein coding domains, and further divided into a number of lineages according to phylogenetic distance (Neumann *et al.*, 2019).

Contrasting with the previous notion that repetitive DNA was no more than “junk DNA”, cytogenomic studies in the last few decades helped to uncover possible roles for tandem and dispersed repeats. Specific satellite DNAs and retrotransposons have been found to have a functional and/or structural role in centromeres (Cheng *et al.*, 2002; Nagaki *et al.*, 2003; Houben & Schubert, 2003; Marques *et al.*, 2015; Macas *et al.*, 2015; Ribeiro *et al.*, 2017). The role of LTR-retrotransposons in genome size variation led to investigations correlating these to heterochromatin distribution, ecological variables, community structure and plant distribution (Guignard *et al.*, 2016; Van-Lume *et al.*, 2017; Lyu *et al.*, 2018; Souza *et al.*, 2019). The activity of transposable elements in the host genome can also cause modifications to gene regulation and the formation of retrogenes, generating morphological innovations and impacting speciation processes (Schrader & Schmitz, 2019). Moreover, lineage-specific satellite DNAs have been widely used as efficient cytomolecular markers, allowing the identification of chromosome pairs and elucidating chromosome rearrangement and duplication events (Koo *et al.*, 2011; Čížková *et al.*, 2013; Ávila Robledillo *et al.*, 2018; Ribeiro *et al.*, 2020).

The fast evolution of repetitive DNA, with many satellite families being genus-or species-specific, impair their use in phylogenetic studies, as concerted evolution and homogenisation reduces sequence variability for comparative studies across taxa (Macas *et al.*, 2015; Mascagni *et al.*, 2020; Ribeiro *et al.*, 2020). Their abundances, however, have phylogenetic signal, as demonstrated by a method to reconstruct phylogenetic relationships using the abundance of different repetitive elements (Dodsworth *et al.*, 2015). This method has proven useful to elucidate relationships in different groups of plants and animals (Dodsworth *et al.*, 2017; Bolsheva *et al.*, 2019; Martín-Peciña *et al.*, 2019). Other methods have assessed the usefulness of repeat-based phylogenetic analysis. More recently, it was demonstrated that sequence similarity measures of repeated sequences can also be used as characters to resolve phylogenetic relationships (Vitales *et al.*, 2019). In a similar approach, assembly and alignment free (AAF) methods can be applied to high complexity fractions of the genome, such as repetitive DNA, in a phylogenomic framework (Fan *et al.*, 2015; Sarmashghi *et al.*, 2019).

In order to identify and characterize the diversity of repeats in a genome, the RepeatExplorer pipeline was created, using a graph-based clustering algorithm to group high-copy sequences based on similarity, with posterior identification of protein-coding domains (Novak *et al.*, 2013). RepeatExplorer can identify repetitive elements with approximately 0.1× genome coverage, most commonly known as genome skimming (GS), which is often sufficient to study high-copy nuclear and organellar DNA (Straub *et al.*, 2012; Dodsworth, 2015; Dodsworth *et al.*, 2019). The GS method is just one of several “reduced representation” methods of high-throughput sequencing. Throughout the last decade, a number of these sequencing methods have been proposed, generating high quality sequencing data at decreasing costs, such as restriction site-associated sequencing (RAD-seq, Eaton et al. 2016) and transcriptome sequencing (RNA-seq, Wang et al. 2017).

Another widely used reduced representation method is target capture sequencing (TCS), in which several genomic probes are designed to “capture” and enrich specific low-copy coding regions of the nuclear genome (Albert *et al.*, 2007; Gnirke *et al.*, 2009). These probes can be designed based on conserved regions retrieved from the alignment of several genomes of a divergent set of organisms (e.g. Anchored Hybrid Enrichment, Lemmon *et al.*, 2012) or by comparing transcriptomic data to search for a conserved set of orthologs across a group (Johnson *et al.*, 2019). Since the development of TCS (Albert *et al.*, 2007), many sets of probes have been developed and applied in both plants and animals (Cosart *et al.*, 2011; Faircloth *et al.*, 2012; Mandel *et al.*, 2014; Ilves & López-Fernández, 2014; Sass *et al.*, 2016; Heyduk *et al.*, 2016; Schmickl *et al.*, 2016). Moreover, the nature of these probe design approaches have allowed the development of universal probe sets, with the potential to be used across all of the angiosperms (e.g. Buddenhagen et al. 2016; Johnson et al. 2019). In addition to the advantage of being universal, high recovery rate of target regions (or the number of enriched targeted loci in the final library) is often achievable with very low “enrichment efficiency” (percentage of library reads successfully mapped to a target sequence) (Johnson *et al.*, 2019), meaning that a TCS enriched library will often contain a high number of “off-target” reads. The use of these off-target reads, in combination with the enriched reads, has been referred to as “Hyb-Seq” (hybrid sequencing, Weitemier *et al.*, 2014). As the off-target reads are often rich in high-copy DNA, Hyb-Seq approaches have been used to reconstruct whole plastomes and ribosomal DNA (Weitemier *et al.*, 2014; Schmickl *et al.*, 2016).

To check whether off-target reads could also be used for identifying, and potentially quantifying, repetitive DNA, we selected the sedge *Rhynchospora* Vahl as a model. *Rhynchospora* is one of the largest genera of Cyperaceae Juss., comprising approximately 350 species, but it is under-studied from a phylogenetic point of view (Thomas *et al.*, 2009; Buddenhagen *et al.*, 2016). Cytologically, *Rhynchospora* has been of great interest due to its holocentric chromosomes, which present the centromere dispersed along the sister chromatids rather than localized in a primary constriction (Bureš *et al.*, 2013). Moreover, cytogenomic studies on *R. pubera* (Vahl) Boeckeler have led to the discovery of the first centromere-specific satellite DNA reported in a holocentric organism, Tyba (Marques *et al.*, 2015). Subsequent studies confirmed the presence of Tyba in other *Rhynchospora* species (Ribeiro *et al.*, 2017). Nevertheless, other non-centromeric satellites found in *Rhynchospora* species showed the typical block-like pattern on localized chromosomal regions (Ribeiro *et al.*, 2017).

Large-scale repeat analysis covering all major clades of *Rhynchospora* would provide valuable insights into the repeat evolution of this genus. Recently, using a set of probes developed by Anchored Hybrid Enrichment (Buddenhagen *et al.*, 2016), a number of *Rhynchospora* species were sequenced using a TCS approach. Here, we assessed the quality of repeat characterization in five *Rhynchospora* species from this dataset. As a considerable part of the off-target reads can be sequences close to the original targeted loci (Dodsworth *et al.*, 2019), we searched for sequencing bias by comparing the target results with results obtained from genome skimming (i.e. unenriched libraries). We specifically addressed three questions: 1) Can we characterize the repetitive DNA fraction of *Rhynchospora* genomes using TCS data, compared to GS data?; 2) Can we develop cytological markers from TCS clustering data?; and 3) Can we use TCS data to reconstruct repeat-based phylogenetic trees?

## MATERIALS AND METHODS

### Plant material and sequence data

Individuals of *Rhynchospora cephalotes* (L.) Vahl and *R. exaltata* Kunth were collected near the towns of Jacaraú (Paraíba, Brazil, voucher UFP87625) and Jaqueira (Pernambuco, Brazil, voucher JPB51537), respectively. These individuals were cultivated in (i) the experimental garden of the Laboratory of Plant Cytogenetic and Evolution at the Universidade Federal de Pernambuco (Brazil), (ii) the greenhouse of the Max Plank Institute for Plant Breeding Research (Cologne, Germany) and (iii) the greenhouse of the Leibniz Institute of Plant Genetics and Crop Plant Research (IPK Gatersleben, Germany).

We downloaded available short read archive data of *R. pubera* (Vahl) Boeckeler (Marques *et al.*, 2015, BioProject PRJEB9643) from the NCBI GenBank (www.ncbi.nlm.nih.gov). Genome skimming sequences of *R. globosa* (Kunth) Roem. & Schult. and *R. tenuis* Link were obtained from Ribeiro *et al*. (2017) and deposited on GenBank under BioProject PRJNA672922. TCS reads (150 bp) for *R. cephalotes, R. exaltata, R. globosa, R. pubera* and *R. tenuis* were obtained from Buddenhagen (2016) and deposited on GenBank under BioProject PRJNA672127.

### DNA extraction and sequencing

Genomic DNA from *R. cephalotes* and *R. exaltata* was extracted from leaves with NucleoBond HMW DNA kit (Macherey and Nagel, Düren, Germany). Quality was assessed with Agilent TapeStation and the gDNA was quantified by Qubit BR assay (Thermo). An Illumina-compatible library was then prepared from 400 ng input gDNA with an NEBNext Ultra™ II FS DNA Library Prep Kit for Illumina (New England Biolabs) with a total of four PCR cycles to introduce dual indexed barcodes. Libraries were sequenced in the Max Planck Genome Centre Cologne on a HiSeq2500 system with 2× 250 bp rapid mode using a HiSeq Rapid PE Cluster and Rapid SBS v2 kit. The new sequence data was deposited on GenBank under BioProject PRJNA672693.

### Genome size measurements

Novel DNA content measurements for *R. cephalotes* and *R. exaltata* were estimated by flow cytometry. Sample preparation was done according to Loureiro *et al.* (2007). Young leaves of each of the studied plants were chopped simultaneously with their respective reference standard, *Solanum lycopersicum* cv. Stupicke (2C = 1.96 pg, Dolezel *et al.*, 1992) for *R. cephalotes* and *Raphanus sativus* L. cv. Saxa (2C = 1.11 pg, Dolezel *et al.*, 1992) for *R. exaltata* in a Petri dish (kept on ice) containing 2 mL of Woody Plant Buffer (WPB). The sample was then filtered through a 30-μm disposable mesh filter (CellTrics, SYSMEX, Norderstedt, Germany) with following addition of 50 μg/mL propidium iodide (from a stock of 1 mg/mL; Sigma-Aldrich) and 50 μg/mL RNase (Sigma-Aldrich). Nine replicates per species were made. The samples were measured in a CyFlow Space flow cytometer (SYSMEX) equipped with a green laser (532 nm). Histograms of relative fluorescence were obtained using the software Flomax v.2.3.0. (SYSMEX, Norderstedt, Germany). Mean fluorescence and coefficient of variation were assessed at half of the fluorescence peak. The absolute DNA content (pg/2C) was calculated multiplying the ratio of the G1 peaks by the genome size of the internal standard.

### Filtering of TCS reads

As our aim was to characterize the repeat fraction of the sequenced species, we were only interested in the off-target reads, whose abundance is inversely proportional to the efficiency of target sequence enrichment. To get rid of the target reads, we filtered the raw TCS data by mapping them to a set of 256 sequences representing consensus sequences of the target loci that were enriched in the *Rhynchospora* dataset (Buddenhagen, 2016), saving the unmapped reads for the repeat characterization. Two different mapping algorithms were used for comparison: the *Geneious read mapper* v6.0.3, with custom sensitivity settings (60% Maximum mismatch per read, Index world length = 12, Maximum ambiguity = 8, Kearse *et al.*, 2012) and the BowTie2 v2.4.1 mapper with high sensitivity settings (0 to 800 insert size, report all matches, Langmead & Salzberg, 2012), both implemented in the software Geneious v.7.1.9 (Kearse *et al.*, 2012). Additionally, we uploaded a FASTA file with the 256 target sequences to RepeatExplorer (https://repeatexplorer-elixir.cerit-sc.cz/) prior to the analysis as a Custom Repeat Database (Novak *et al.*, 2013). With this option we could exclude clusters of enriched gene sequences mistakenly identified as repeats from the analysis. In summary, we ended with one GS dataset (comprising of previously published datasets for *R. globosa, R. pubera* and *R. tenuis* and the newly sequenced *R. cephalotes* and *R. exaltata* data) and four different “target datasets” for each species: 1) Raw target capture reads; 2) Reads left after mapping with *Geneious read mapper*; 3) Reads left after mapping with BowTie Mapper and 4) Reads left after exclusion of “enriched gene clusters” annotated according to a custom Repeat Database (RE).

### In silico Repeat Analysis

In order to compare the repeat composition observed in all different datasets, we employed the RepeatExplorer pipeline (Novak *et al.*, 2013). Reads from all datasets of *R. cephalotes, R. exaltata, R. globosa, R. pubera* and *R. tenuis* were uploaded to the platform, filtered by quality with default settings (95% of bases equal to or above the quality cut-off value of 10) and interlaced. Clustering was performed with default settings of 90% similarity over a 55% minimum sequence overlap. The *Find RT Domains* tool and additional database searches (Genbank) were used to identify protein domains for repeat annotation, and graph layouts of individual clusters were examined interactively using the SeqGrapheR tool (Novak *et al.*, 2013).

Although the entire set of reads from each dataset was uploaded to RepeatExplorer, we used the *Read Sampling* option on the clustering analysis to manually input the number of reads to be analysed, accounting for 0.13× of the genome of each species (Table 1). Only for *R. globosa* was this not possible, as no information on genome size was available for this species. In this case, we ran tests with 500,000, 1,000,000 and 2,000,000 reads. However, independently of the number of reads used as input, only 200,061 reads were analysed, always with similar results. Clusters with at least 0.01% genome abundance were automatically annotated and manually checked. The genome proportion of high copy repeats was estimated based on the number of reads of the annotated clusters.

**Table 1-.** Genome size (2C), number of reads from GS (genome skimming) and raw TCS datasets, percentage of mapped reads on Bowtie Mapper and Geneious Read Mapper datasets, percentage of reads identified on “target clusters” of RepeatExplorer (RE) dataset and number of reads analysed for all datasets on each species.

| Species | 1C<br>(Mbp) | Number of<br>GS read<br>pairs | Number of<br>raw TCS<br>reads<sup>d</sup> | % of filtered TCS reads* |  |  | Read pairs<br>analysed |
| --- | --- | --- | --- | --- | --- | --- | --- |
|  |  |  |  | BowTie | Geneious | RE |  |
| *Rhynchospora cephalotes*<br>(L.) Vahl | 356.97 | 11,737,291 | 6,932,932 | 11.77 | 23.12 | 16.80 | 600,000 |
| *Rhynchospora exaltata*<br>Kunth | 244.5 | 21,384,261 | 5,151,874 | 13.02 | 20.65 | 18.45 | 422,199 |
| *Rhynchospora globosa*<br>(Kunth) Roem. & Schult. | - | 4,000,000<sup>c</sup> | 3,713,422 | 4.10 | 14.95 | 8.82 | 200,061 |
| *Rhynchospora pubera*<br>(Vahl) Boeckeler | 1,613.7<sup>a</sup> | 4,000,000<sup>a</sup> | 4,164,982 | 3.96 | 12.29 | 8.66 | 2,797,919 |
| *Rhynchospora tenuis*<br>Link | 381.42<sup>b</sup> | 4,000,000<sup>c</sup> | 4,826,084 | 10.18 | 20.64 | 6.36 | 661,128 |

<sup>a</sup>Marques *et al.*, 2015; <sup>b</sup>Ribeiro *et al.*, 2018; <sup>c</sup>Ribeiro *et al.*, 2017; <sup>d</sup>Buddenhagen 2016
\*% of target reads identified by the different strategies

We used the TAREAN tool (Novák *et al.*, 2017) available in the RepeatExplorer pipeline to annotate satellite DNAs. Satellites were named based on previous publications or by using the species abbreviation followed by SAT, a number based on the decreasing order of abundance and a hyphen followed by the number of basepairs of the monomer. The consensus monomer sequences of the identified satellite DNAs of each species were compared using DOTTER (Sonnhammer & Durbin, 1995).

### Testing Method Performance

To assess if the abundances of the different repeats observed in our raw and filtered target capture datasets were comparable, we compared their abundances with the ones observed in the GS datasets. For this, we used the abundance values of the individual repeat families (Table S1). As the data did not follow a normal distribution, we used Spearman’s rank correlation. The analysis was undertaken with the package *stats* implemented in the software R v4.0.2 (R Core Team, 2019). Correlation plots were constructed with the R package ggplot2 (Wickham, 2016).

### Repeat amplification, probe labelling and *in situ* hybridization

Primers for the *R. cephalotes* Tyba variant found in the TCS datasets (see results) were designed based on the most conserved region of the consensus sequence (F: 5’-AAGCTATTTGAATGCAATTATGTGC; and R: 5’AGCGTTTCTAGCCACATTTGA). Genomic DNA (40 ng) of *R. cephalotes* was used for PCR reaction with 1× PCR buffer, 2 mM MgCl_2_, 0.1 mM of each dNTP, 0.4 μM of each primer, 0.025 U Taq polymerase (Qiagen) and water. The PCR conditions were as follow: 94°C 2 min, 30 cycles of 94°C 50 s, 58°C 50 s and 72°C 1 min and 72°C 10 min. PCR products were labelled with Atto488-dUTP (Jena Bioscience) with a nick translation labelling kit (Jena Bioscience).

Mitotic chromosomes of *R. cephalotes* were prepared from root tips, pre-treated in 2 mM 8-hydroxyquinoline at 10°C for 20 h and fixed in ethanol: acetic acid (3:1 v/v) for 2 h at room temperature and stored at −20°C. Fixed root tips were digested with 2% cellulose, 2% pectinase and 2% pectolyase in citrate buffer (0.01 M sodium citrate and 0.01 M citric acid) for 120 min at 37°C and squashed in a drop of 45% acetic acid. Fluorescent *in situ* hybridization was performed as described by Aliyeva-Schnorr *et al.*, (2015). The hybridization mixture contained 50% (v/v) formamide, 10% (w/v) dextran sulfate, 2×SSC, and 5 ng/μl of the probe. Slides were denatured at 75°C for 5 min, and the final stringency of hybridization was 76%.

### Immuno-FISH

To visualize the centromeres of *R. cephalotes*, we performed immunostaining of the centromere-specific histone variant CENH3 with polyclonal antibodies developed for *R. pubera* (RpCENH3, Marques *et al.*, 2015). Mitotic preparations were made from root meristems fixed in 4% paraformaldehyde in Tris buffer (10 mM Tris, 10 mM EDTA, 100 mM NaCl, 0.1% Triton, pH 7.5) for 5 minutes on ice under vacuum and for another 25 minutes only on ice. After washing twice in Tris buffer, the roots were chopped in LB01 lysis buffer (15 mM Tris, 2 mM Na_2_EDTA, 0.5 mM spermine 4HCl, 80 mM KCl, 20 mM NaCl, 15 mM β-mercaptoethanol, 0.1% Triton X-100, pH 7.5), filtered through a 50 μm filter (CellTrics, Sysmex), diluted 1:10, and subsequently, 100 μl of the diluted suspension were centrifuged onto microscopic slides using a Cytospin3 (Shandon, Germany) as described by Jasencakova *et al.* (2001). Immuno-FISH with anti-RpCENH3 antibodies and the Tyba repeat was performed according to Ishii *et al.* (2015). We used rabbit anti-RpCENH3 (diluted 1:200) as primary antibody and detected it with Cy3-conjugated anti-rabbit IgG (Dianova) secondary antibody (diluted 1:200). Slides were incubated overnight at 4°C and washed three times in 1×PBS before the secondary antibody were applied.

### Microscopy

For widefield microscopy, we used an epifluorescence microscope BX61 (Olympus) equipped with a cooled CCD camera (Orca ER, Hamamatsu). To achieve super-resolution of ~120 nm (with a 488 nm laser excitation), we applied spatial structured illumination microscopy (3D-SIM) using a 63×/1.40 Oil Plan-Apochromat objective of an Elyra PS.1 microscope system and the software ZENBlack from Carl Zeiss GmbH (Weisshart *et al.*, 2016).

### Comparative repeat phylogenomics

We employed a repeat abundance-based phylogenetic inference method (see details in Dodsworth *et al.*, 2015) to assess if repeat abundance identified in our TCS reads could be used to resolve phylogenetic relationships, using one of our filtered datasets (BowTie) and the GS dataset for comparison. First, we concatenated reads for our five species with 0.065× coverage, with species-specific codes for each set of reads, and ran a comparative clustering analysis (simultaneous clustering of all species on the dataset) on RepeatExplorer with default settings (Novak *et al.*, 2013). As the sequences were coded with the species names, we could identify the number of reads that each species contributed to each of the generated clusters, which is proportional to genome. Parsimony analysis using repeat abundances as quantitative characters was undertaken as described by Dodsworth et al. (2015).

To access the phylogenetic potential of repetitive elements based on sequence similarity, we used the alignment and assembly free (AAF) approach (Fan *et al.*, 2015) using all reads identified as repeats by RepeatExplorer in the Bowtie dataset. AAF constructs phylogenies directly from unassembled genome sequence data, bypassing both genome assembly and alignment. Thus, it calculates the statistical properties of the pairwise distances between genomes, allowing it to optimize parameter selection and to perform bootstrapping.

In order to compare our repeat abundance-based phylogeny with a nuclear marker-based phylogeny, we extracted the aligned sequences of 256 loci gathered by target capture (Buddenhagen, 2016) of our five *Rhynchospora* species. For simplification, we used the most general model of DNA substitution GTR + I + G (Abadi *et al.*, 2019). Phylogenetic relationships were inferred using Bayesian Inference (BI) as implemented on BEAST v.1.8.3 (Drummond & Rambaut, 2007). Two independent runs with four Markov Chain Monte Carlo (MCMC) were conducted, sampling every 1,000 generations for 10,000,000 generations. Each run was evaluated in TRACER v.1.7 (Rambaut *et al.*, 2018) to assess MCMC convergence and a burn-in of 25% was applied. We then obtained the consensus phylogeny and clade posterior probabilities with the “sumt” command.

## RESULTS

### Efficiency of target sequences filtering

We generated GS data and genome size estimates for *Rhynchospora cephalotes* and *R. exaltata* to add to the already sequenced data of the other three *Rhynchospora* here analysed in order to compare repeat composition from TCS and GS data (Table 1). The Geneious datasets presented a higher number of filtered reads than the BowTie datasets (Table 1). *R. pubera* presented the smallest number of filtered reads among all five species, with both the BowTie and Geneious filters (3.96% and 12.29% respectively). *R. cephalotes* had the highest number of filtered reads among the Geneious datasets, while *R. exaltata* presented the largest proportion among the BowTie datasets. Using the Custom Repeat Database option of RepeatExplorer, the highest amount of “target clusters” was found on *R. exaltata* (18.45%) and the lowest amount was found on *R. tenuis* (6.36%).

### Repetitive DNA content of different datasets

To evaluate the quality of repeat characterization from TCS datasets, the genomic proportion of different repetitive element lineages was compared with those observed in the GS datasets (Table 2, Fig. 1). Generally, the proportion of the total repeat fraction observed in the GS datasets was smaller than the ones observed in all the TCS datasets (Table 2, Figure S1). The raw datasets presented the highest values for total repetitive fraction in almost all species, with the exception of *R. globosa*, in which the Geneious dataset showed a total of 49.41% repeat proportion against 46.74% on the raw dataset. This is also reflected in the difference of number of clusters representing at least 0.01% of total genomic proportion formed in the clustering analysis. While the GS datasets presented cluster numbers varying from 155 to 294, filtered TCS datasets ranged from 462 to 589 clusters and raw TCS datasets ranged from 628 to 751. These differences are mostly due to the discrepancy in the proportion of unclassified repetitive elements in the different datasets (Table 2, Fig. S1). Unclassified repeat proportion on GS varied from 6.21% in *R. exaltata* to 14.70% in *R. globosa*, while in TCS it accounted for to up to 43.63% (raw TCS of *R. exaltata).* Overall, proportion of unclassified repeats was smaller in all filtered datasets when compared to raw TCS (Fig. 2a). Additional mapping to the original 256 target regions and separated BLAST searches did not produce any matches for the unclassified clusters, suggesting that these could be either repetitive or non-repetitive accidentally enriched genomic regions, too fragmented to be annotated.

**Table 2-.** Genomic abundance of repeat types for all datasets in the five analysed *Rhynchospora* species

| Species | Dataset | Total Repeat fraction | Unclassified Repeats | 5S rDNA | 35S rDNA | Satellites | Class I/ Unc. LTRs | Class I/ LTR Ty1/copia | Class I/ LTR Ty3/gypsy | Class I/ Pararetrovirus | Class I/ LINE | Class II/ Subclass 1 | Class II/ Subclass 2 |
| --- | --- | --- | --- | --- | --- | --- | --- | --- | --- | --- | --- | --- | --- |
| *R. cephalotes* | GS | 14.81 | 7.39 | 0.06 | 0.40 | 3.76 | 0.66 | 0.69 | 0.95 | 0.29 | 0 | 0.58 | 0.03 |
|  | Raw | 43.08 | 38.45 | 0.11 | 0.21 | 2.84 | 0.51 | 0.16 | 0.45 | 0.05 | 0.06 | 0.24 | 0 |
|  | Geneious | 33.38 | 28.54 | 0.15 | 0.33 | 2.58 | 0.77 | 0.29 | 0.32 | 0.08 | 0 | 0.32 | 0 |
|  | BowTie | 34.75 | 30.29 | 0.13 | 0.30 | 2.52 | 0.37 | 0.27 | 0.47 | 0.13 | 0.01 | 0.26 | 0 |
|  | RE | 31.17 | 25.92 | 0.13 | 0.25 | 3.44 | 0.61 | 0.19 | 0.29 | 0.05 | 0 | 0.29 | 0 |
| *R. globosa* | GS | 38.91 | 14.70 | 0.04 | 0.45 | 4.19 | 3.25 | 1.93 | 12.51 | 0.16 | 0.01 | 1.67 | 0 |
|  | Raw | 46.74 | 27.05 | 0.11 | 0.76 | 1.63 | 0.84 | 2.68 | 12.12 | 0.17 | 0.12 | 1.26 | 0 |
|  | Geneious | 49.42 | 25.20 | 0.10 | 0.79 | 1.86 | 1.08 | 2.96 | 15.50 | 0.20 | 0.13 | 1.60 | 0 |
|  | BowTie | 45.59 | 23.84 | 0.12 | 0.69 | 1.74 | 0.39 | 2.89 | 14.25 | 0.18 | 0.11 | 1.38 | 0 |
|  | RE | 42.13 | 20.52 | 0.12 | 0.84 | 1.79 | 0.93 | 2.94 | 13.29 | 0.19 | 0.13 | 1.38 | 0 |
| *R. tenuis* | GS | 17.99 | 8.23 | 0.02 | 0.38 | 2.61 | 0.37 | 5.64 | 0.47 | 0 | 0.03 | 0.24 | 0 |
|  | Raw | 27.35 | 19.53 | 0.08 | 0.58 | 3.06 | 0.54 | 1.96 | 0.74 | 0.04 | 0.08 | 0.74 | 0 |
|  | Geneious | 24.41 | 15.59 | 0.05 | 0.68 | 3.45 | 0.71 | 2.22 | 0.77 | 0.04 | 0.07 | 0.83 | 0 |
|  | BowTie | 23.79 | 15.36 | 0.08 | 0.69 | 3.19 | 0.66 | 2.13 | 0.71 | 0.04 | 0.05 | 0.88 | 0 |
|  | RE | 22.37 | 14.13 | 0.08 | 0.62 | 3.28 | 0.54 | 2.00 | 0.80 | 0.04 | 0.09 | 0.79 | 0 |
| *R. exaltata* | GS | 27.25 | 6.21 | 0.10 | 2.14 | 16.10 | 0 | 2.14 | 0.44 | 0 | 0.07 | 0.05 | 0 |
|  | Raw | 49.87 | 43.63 | 0.10 | 1.10 | 0.91 | 0.57 | 1.43 | 1.72 | 0 | 0.13 | 0.28 | 0 |
|  | Geneious | 44.28 | 33.53 | 0.16 | 1.31 | 3.69 | 0.67 | 2.01 | 2.13 | 0.02 | 0.11 | 0.65 | 0 |
|  | BowTie | 43.67 | 33.96 | 0.11 | 1.04 | 3.39 | 0.60 | 2.24 | 1.63 | 0 | 0.09 | 0.61 | 0 |
|  | RE | 41.58 | 31.53 | 0.12 | 1.20 | 3.48 | 0.65 | 1.94 | 1.99 | 0.01 | 0.15 | 0.51 | 0 |
| *R. pubera* | GS | 26.67 | 9.87 | 0 | 0.61 | 3.87 | 2.12 | 6.78 | 0.85 | 0.46 | 0.22 | 1.89 | 0 |
|  | Raw | 39.11 | 23.48 | 0 | 1.82 | 1.04 | 0.48 | 6.74 | 4.08 | 0.46 | 0.51 | 0.43 | 0.07 |
|  | Geneious | 39.11 | 19.09 | 0 | 1.99 | 1.17 | 0.67 | 7.53 | 6.51 | 0.55 | 0.59 | 1.01 | 0 |
|  | BowTie | 36.92 | 20.44 | 0 | 2.04 | 0.91 | 0.31 | 7.26 | 3.81 | 0.50 | 0.55 | 1.1 | 0 |
|  | RE | 38.31 | 18.66 | 0 | 2.82 | 0.33 | 0.51 | 7.23 | 6.62 | 0.49 | 0.54 | 1.04 | 0.07 |

**Figure 1 –.**
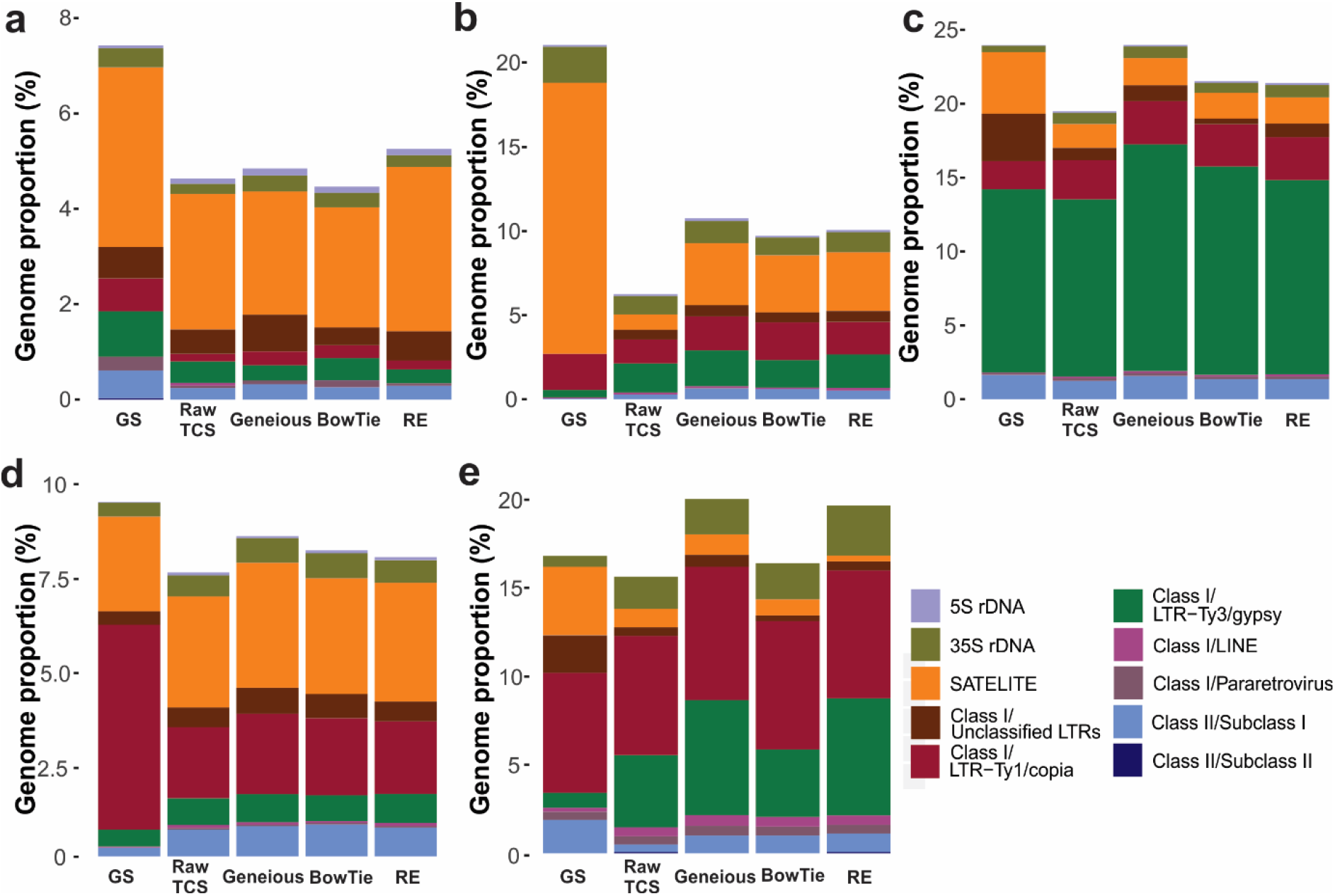
Barplots representing genomic abundance of classified repeat types identified in every dataset of *Rhynchospora cephalotes* (a), *R. exaltata* (b), *R. globosa* (c), *R. pubera* (d) and *R. tenuis* (e). Bar colours represent different repeat types according to the caption at the lower right corner.

**Figure 2 –.**
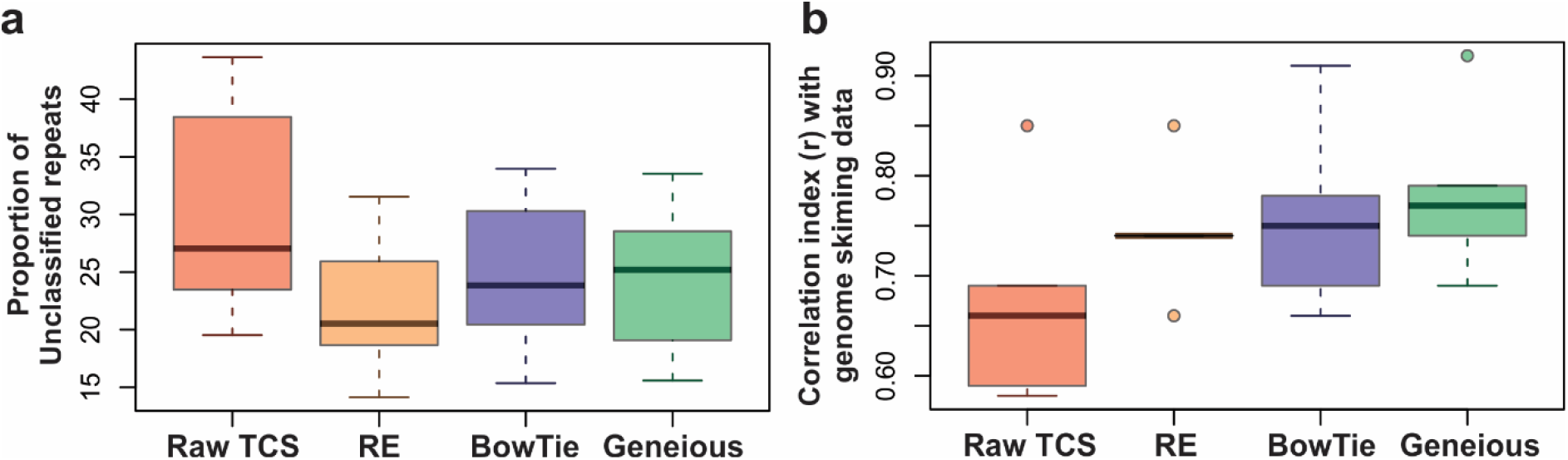
Comparison of genomic proportions of unclassified elements (a) and correlation index (r) with genome skimming data (b) between raw and filtered Target Capture datasets (RepeatExplorer, BowTie and Geneious) of all *Rhynchospora* species.

The filtering strategies did not largely impact satellite DNA abundance in the analysed species, with the exception of *R. exaltata*, in which it was possible to identify ~4× more satellite reads in the BowTie and Geneious datasets than in the raw datasets. Also in *R. exaltata*, there was a huge discrepancy between the satellite abundance observed in the GS dataset when compared to all TCS datasets (Fig. 1). However, the two satellite DNAs responsible for this abundance difference were also found in all TCS datasets (Table S2, Fig. S2), although in smaller proportions. The amount of satellite DNA was generally lower in the TCS datasets, with the exception of *R. tenuis*, in which all TCS datasets presented a small increase in satellite abundance when compared to GS (Table 2, Fig. 1). Although in some species some of the satellites found in GS data were not present in every dataset, the most abundant for each species could be identified in all of the TCS datasets. Similarly, some satellites found in TCS datasets were not found in the GS datasets, possibly being low-abundance satellites accidentally enriched (Table S2, Fig. S2).

For mobile elements, there was a general agreement in the order of abundance of repeat types found in the GS and TCS datasets, with filtered datasets showing increase in the proportion of annotated elements when compared to the raw TCS dataset (Table 2, Fig. 1), with a few exceptions. For example, in *R. exaltata*, LTRs from the Ty3/gypsy superfamily were more abundant in the raw TCS, BowTie and RE datasets than in the GS datasets (Table 2, Fig. 1). We also compared the abundances at lineage level (Table S1). Patterns of abundance of LTR families from GS and TCS datasets were similar, with most of the genomic abundance of Ty1/copia and Ty3/gypsy superfamilies being result of the amplification of up to four main lineages (Table S1). Generally, LTRs found in GS with abundance as low as 0.01% could also be identified in the target datasets. Surprisingly, in all five species, a greater diversity of LTR retroelements was observed in the target capture datasets when compared to GS (Table S1). This led to a few interesting discrepancies, such as in *R. pubera*, where target capture datasets showed high abundance of Ty3/gypsy/Retand (1 to 1.3%) and Ty3/gypsy/Tekay elements (0.46 to 0.50%), despite those not being found in GS data.

To statistically compare the results obtained in the different datasets, we checked for a correlation between the classified repeat abundances observed (at lineage level, when possible) in all target capture datasets with the ones observed in the GS datasets (Fig. 3). Although all tests showed significant correlations (*p* < 0.05), the strength of the correlation varied depending on the dataset. Raw TCS datasets had the weakest correlation in all five species, with filtering of the targeted sequences generally improving the correlation with the GS dataset (Fig. 2b). The strongest correlations for *R. globosa* (*r* = 0.92), *R. pubera* (*r* = 0.74) and *R. tenuis (r* = 0.79) were observed with the Geneious dataset, while for *R. cephalotes* and *R. exaltata* the best correlation was observed on the BowTie dataset (*r* = 0.78 and 0.75). The RE filtering for *R. globosa* and *R. tenuis* did not improve the correlation when compared to the raw TCS dataset (*r* = 0.85 and *r* = 0.66 respectively). However, it also showed as strong a correlation as BowTie for *R. cephalotes* (*r* = 0.78) and as Geneious for *R. pubera* (*r* = 0.74).

**Figure 3 –.**
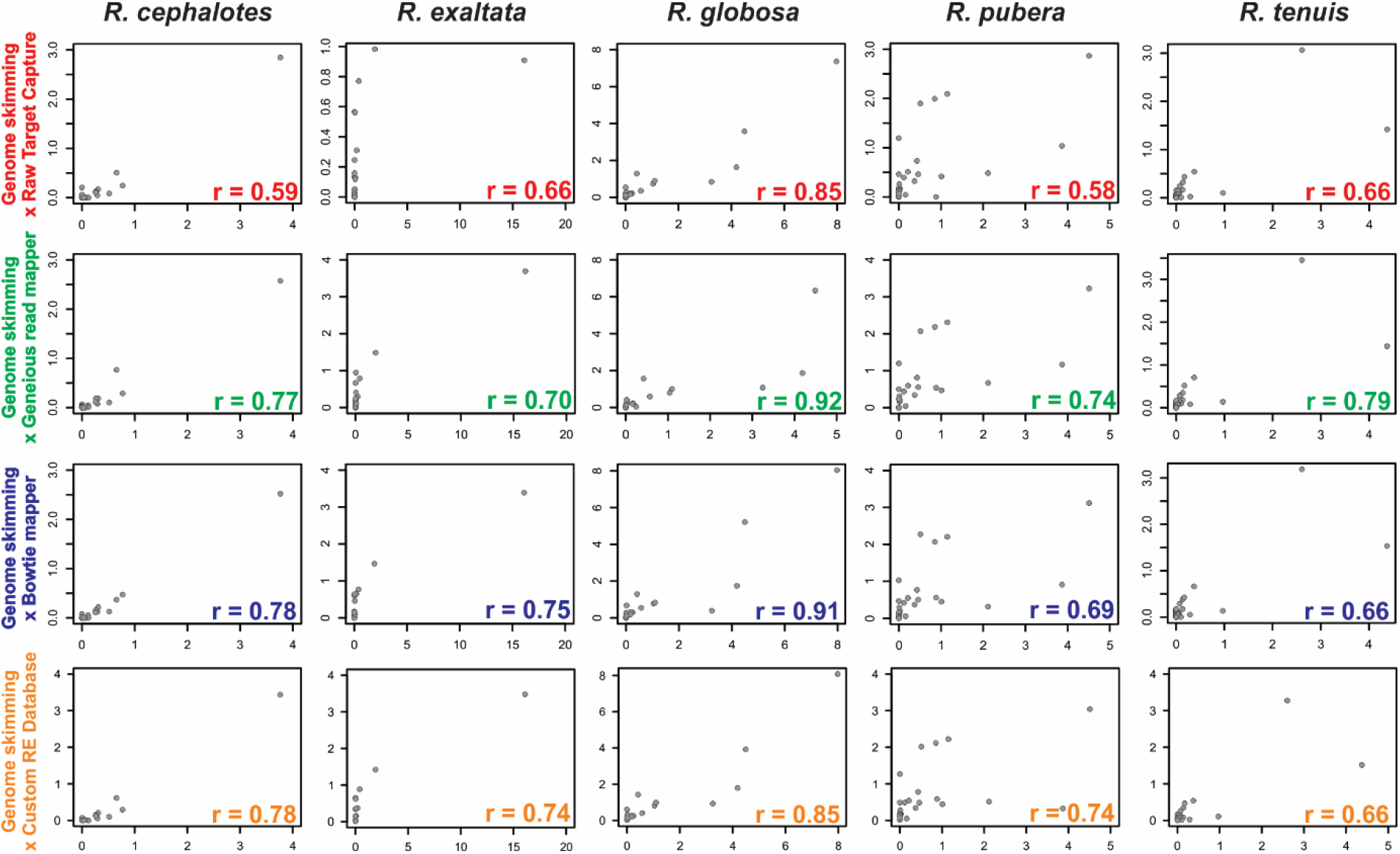
Correlation between repeat abundances observed on genome skimming and target capture datasets of *Rhynchospora* species. Genome skimming abundance values are represented on the x axis of each plot, while target dataset abundances are on the y axis. Spearman’s correlation index (r) for each case is plotted in the lower right corner.

### Chromosomal localization of the satellite DNA found in the TCS dataset

To test whether it was possible to use the TCS data to investigate the chromosomal repeat distribution by FISH, we chose the most abundant repetitive element found in the *R. cephalotes* TCS datasets. This repeat was a 172-bp satellite DNA with 60% sequence similarity to Tyba of *R. pubera* (Marques *et al.*, 2015). The *in situ* hybridization pattern of the *R. cephalotes* Tyba (RcTyba) variant was similar to the distribution reported in *R. pubera*. Small foci appeared in interphase nuclei, and a continuous line along both condensed chromatids occurred at all pro- and metaphase chromosomes. Via immuno-FISH using a CENH3-specific antibody and RcTyba repeats, respectively, the holocentric centromere structure of *R. cephalotes* has been confirmed due to the co-localization of CENH3 and RcTyba (Fig. 4).

**Figure 4 –.**
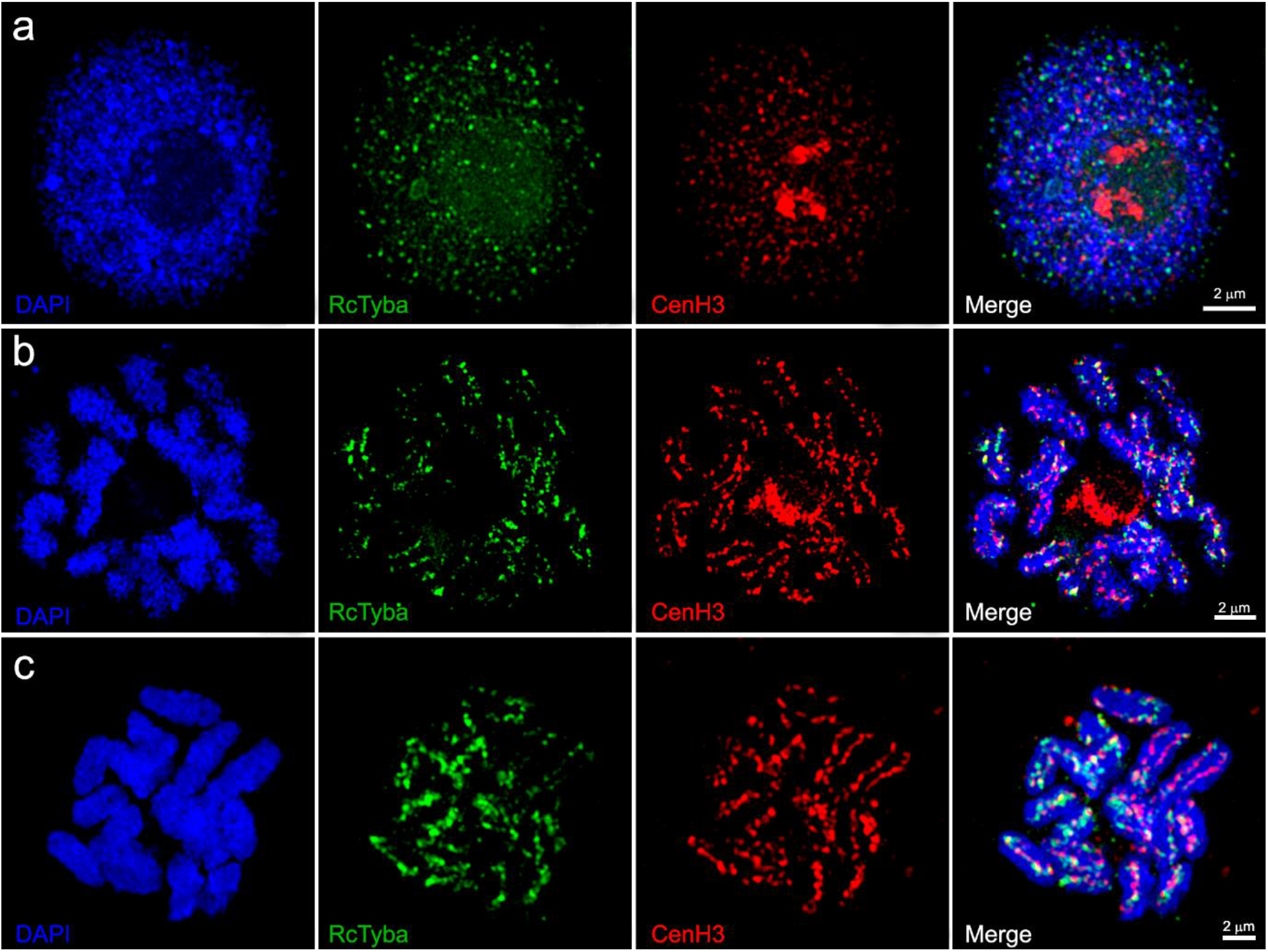
Co-localization of *R. cephalotes* Tyba repeats and CENH3 in interphase nuclei (a), prometaphase (b) and metaphase chromosomes (c) via 3D-SIM imaging. Both Tyba and CENH3 clearly indicate the presence of holocentromeres in the condensed mitotic chromosomes.

### Repeat abundance and structure found in TCS data reflect phylogenetic relationships

We used repeat abundances obtained by comparative clustering analysis of BowTie datasets of our five *Rhynchospora* species to reconstruct phylogenetic relationships. In the comparative clustering analysis of the GS dataset, 1,754,326 concatenated reads were analysed, forming 450 clusters with at least 0.01% genomic abundance. For the BowTie dataset, 1,186,605 of concatenated reads were analysed, with 582 clusters representing at least 0.01% of total genomic abundance. Repeat composition varied among species, with the largest clusters of each species being almost or completely absent in the others (Fig. 5a). By using the first 150 most abundant clusters of the comparative analysis, we were able to reconstruct the phylogenetic relationships among the five *Rhynchospora* species for both the BowTie (Fig. 5b) and GS datasets (Fig. 5c) with high bootstrap support (mean BS = 100 and 98.3 respectively). Using the reads from all clusters identified in the BowTie dataset comparative analysis, the AAF analysis yielded the same relationships with high bootstrap support (mean BS = 100, Fig. 4d). Branch lengths of the abundance-based analysis were significantly higher than for the AAF analysis. Despite this, the species relationships observed in the repeat-based phylogenies were congruent with the ones retrieved in the Bayesian analysis with 256 concatenated target loci, with *R. cephalotes* + *R. exaltata* forming a clade sister to *R. pubera* + *R. tenuis*, and *R. globosa* sister to both clades (Fig. 5e).

**Fig. 5 –.**
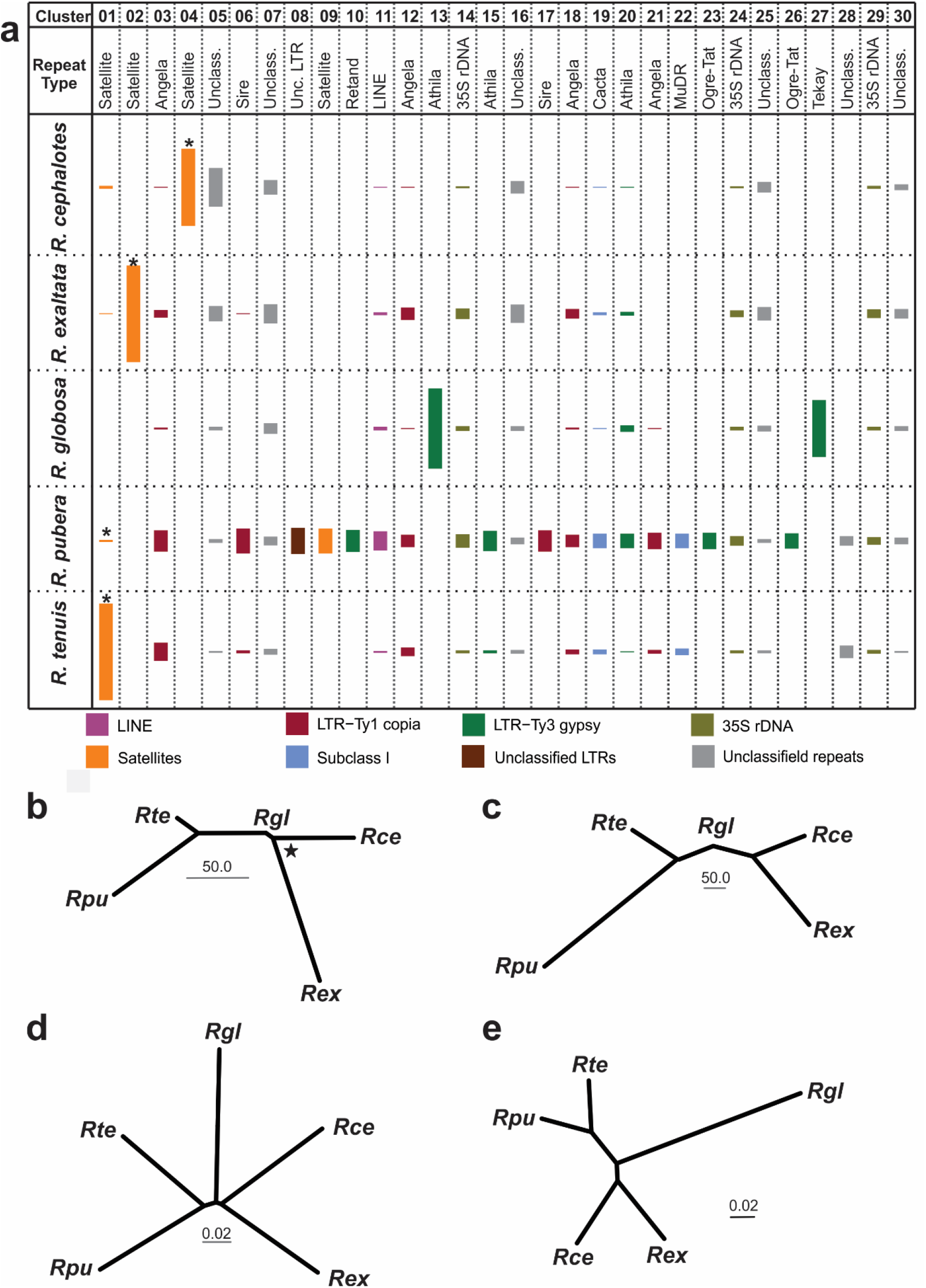
Phylogenomics of *Rhynchospora* species using genome skimming and target capture datasets a) Graphic representation of the 30 most abundant clusters originated from the comparative clustering analysis with the BowTie dataset. Height of rectangles represent the genomic abundance of each cluster, with colours indicating the repeat type according to the caption. Asterisks (*) indicate the clusters that represent Tyba, which is absent in *Rhynchospora globosa*. Below, phylogenetic trees obtained by repeat abundance of the GS (b) and BowTie (c) datasets, AAF of total reads in repetitive clusters (d) and Bayesian inference based on 256 target regions (e). Star in b represents the only node with support below 100 (BS = 99).

## DISCUSSION

### TCS data can be used to characterize the repeat composition of a genome

We were able to find most of the repeat diversity of five *Rhynchospora* species using filtered and unfiltered TCS reads. Depending on the filtering strategy employed, the *Rhynchospora* data used here showed low abundance of on-target reads, indicating a considerable proportion of off-target reads suitable for repeat analysis. Target-based sequencing approaches often have varied enrichment efficiency, with on-target enriched reads representing as low as 5% of the final genomic library (Johnson *et al.*, 2019). This is particularly the case with universal kits that are designed to work across large taxonomic groups, at the cost of enrichment efficiency. Thus, the off-target sequence reads from such libraries can be used in a similar fashion to genome skimming sequencing, an approach often described as Hyb-Seq (Weitemier *et al.*, 2014; Dodsworth *et al.*, 2019).

Our results show that there is a moderate rank correlation between the abundances of annotated repeats obtained by analysing genome skimming and raw target capture datasets, and that this correlation increases when analysing off-target reads only. The significant correlation indexes showed that although annotated repeat proportions of our filtered target datasets are not identical to those observed in GS, repeat composition and order of abundance was highly similar. Although the Geneious dataset presented the highest proportion of filtered reads and highest correlation index with GS data for three of the five species, the BowTie and RE approaches were also sufficient for increasing the correlation index in most cases. Therefore, any of the filtering strategies presented here can produce sufficiently accurate results in regards to repeat identification and order of abundance when compared with the traditional GS approach. It is important to note that, although the patterns of repeat abundance are highly similar, the numerical difference observed for some repeat classes shows that combining GS and TCS data may produce inconsistent results, and exact quantification from these data should be taken with caution.

Although the total abundance of satellite DNAs varied between GS and TCS datasets, we were able to identify the most abundant satellite DNA families of all five *Rhynchospora* species in all target datasets. While some low-abundance satellites (<0.2% of genomic abundance) found in GS data were not found in the target datasets, others with abundances as low as 0.09% were found in all datasets, suggesting that satellite abundance does not necessarily affect its presence in the off-target reads (Table S2). More importantly, various satellites found in the TCS datasets were not found in the GS dataset (Table S2), probably being low-abundant satellites accidentally enriched in the sequencing process. Low-abundant satellites may sometimes not be detected by RepeatExplorer, requiring additional filtering to be detected (Ruiz-Ruano *et al.*, 2016), which could explain the absence of these satellites in our GS datasets.

One of the benefits of using *Rhynchospora* as a model for this study was the fact that it possesses satellite DNAs with varying chromosomal distributions. Tyba clusters are dispersed, forming a linear distribution along the metaphase holocentromeres of *R. pubera*, *R. ciliata*, *R. cephalotes* and *R. tenuis* (Marques *et al.*, 2015; Ribeiro *et al.*, 2017; this study). However, other satellites in *Rhynchospora*, such as *Rg*SAT1-186 in *R. globosa* form localized blocks (Ribeiro *et al.*, 2017). Off-target sequences used in Hyb-Seq are often adjacent to the targeted gene regions (Weitemier *et al.*, 2014; Dodsworth *et al.*, 2019), rasing the possibility that our analysis would preferentially identify widespread repeats such as Tyba, which are interspersed with genic regions (Marques *et al.*, 2015). However, *Rg*SAT1-186, identified as sub-terminal clusters in the chromosomes of *R. globosa* (Ribeiro *et al.*, 2017), was identified as the most abundant cluster on all the *R. globosa* TCS datasets, showing that the chromosomal distribution of a satellite DNA was not interfering in the randomness of the off-target reads (Fig. S1).

As well as confirming its presence in the target datasets of *R. pubera* and *R. tenuis*, we were able to find novel Tyba variants in *R. cephalotes* and in *R. exaltata*, with 60% and 57% sequence similarity to *Rp*Tyba, respectively. *R. globosa* did not present any Tyba variant, in concordance with the results of Ribeiro *et al.* (2017). The abundance of Tyba in *R. pubera* was very low in the target datasets (~0.14%) compared to the GS results (~2.8% here, 3.6% in Marques *et al.* 2015). In our *R. pubera* TCS datasets, the most abundant tandem repeat was *Rp*SAT5-287, which appeared as only the fifth most abundant in the GS dataset (Table S2). This satellite was not found in the previous *R. pubera* characterization and may indicate the potential to discover additional low-abundant sequences using TCS datasets. Although fast rates of evolution for satellite DNAs may lead to intraspecific abundance variation (Ceccarelli *et al.*, 2011), this 20-fold difference is probably too high to be a product of differences between the individuals used for each sequencing method. Satellite abundances estimated by TAREAN depend on several factors, such as sequence coverage, monomer homogeneity and similarity with other repeats (Novák *et al.*, 2017). As abundance information gathered by short reads can grossly underestimate the true abundance of this repeat type (i.e. Ribeiro *et al.*, 2020), caution is needed when interpreting TCS-yielded genomic abundances, though similar caution is needed with GS data as well.

Transposable elements, particularly LTR-retrotransposons, are often the largest fraction of repetitive DNA in plants (Galindo-González *et al.*, 2017). In our GS datasets, LTR-Ty1/copia was the most abundant repeat type in three out of five species. The TCS datasets yielded similar results to GS, with a few discrepancies, especially in *R. exaltata* (predominance of Ty3/gypsy instead of Ty1/copia) and *R. pubera* (significantly higher proportion of Ty3/gypsy than in GS). In order to do a more detailed comparison, we used the individual lineage abundance values for our correlation analysis. The high correlation rates indicate that different LTR lineages, although varying in abundance, contributed similarly to the repetitive fraction in GS and filtered target datasets. Our results show that in addition to being able to find the majority of amplified lineages of LTR retrotransposons, we could also find lineages with abundances as low as 0.01% in the target datasets. We also find a higher diversity of LTR lineages in the target datasets than in the GS data, with a few of these “extra” lineages being over-abundant when compared to the GS dataset (Table S1). These lineages were absent in the GS results probably due to masking by the highly abundant satellite DNA clusters, which were underestimated in some of our TCS datasets. Nonetheless, the fact that we could identify even low abundance LTRs in the target datasets, coupled with the moderate correlation with the GS-yielded abundances, indicate that off-target reads from target sequencing may be sufficient to identify most of the LTR retrotransposons in a genome.

### TCS data can be used to develop cytogenetic probes

We tested whether we could use repeat information obtained from TCS data to develop probes for cytogenetic techniques such as fluorescent *in situ* hybridization (FISH). Highly abundant repetitive elements are frequently chosen for FISH experiments, as they are easier to visualize on chromosomes than low abundance repeats and can often be important components of predominantly heterochromatic and centromeric regions (Marques *et al.*, 2015 Bilinski *et al.*, 2017; Ávila Robledillo *et al.*, 2018). In our *Rhynchospora* species, the most abundant cluster in all datasets was a satellite DNA. For *R. tenuis* and *R. globosa* we found the same satellites found previously by Ribeiro *et al.* (2017) as the most abundant satellites (Tyba and *Rg*SAT1-186 respectively).

Similar to previous results on other *Rhynchospora* (Marques *et al.*, 2015; Ribeiro *et al.*, 2017), the Tyba variant from *R. cephalotes* presented a dispersed distribution in interphase nuclei and a line-like pattern along the sister chromatids of condensed chromosomes. Co-localization with CENH3 further confirmed the holocentromere specific localization of Tyba. The conservation of the holocentromeric distribution of *Rc*Tyba on *R. cephalotes* strengthens its putative role in centromere function as proposed previously (Marques *et al.*, 2015; Ribeiro *et al.*, 2017). It also points to a remarkably old origin for this satellite and its holocentromeric association, sharing a common ancestor between 35-46 My (95% credible interval; Buddenhagen, 2016). A large-scale survey of Tyba in the entire *Rhynchospora* genus could help to further elucidate the evolution of this satellite and its association with the holocentromere. In light of our results, we believe that using off-target reads from target capture-based phylogenetic studies could be an additional way to find potentially informative cytological markers across a broader sampling and investigate their evolution in a broader context.

### TCS data can be used to construct repeat-based phylogenies

In order to test if repeat abundances observed in target datasets are accurate enough to reconstruct phylogenetic relationships, we applied the methodology described by Dodsworth *et al.* (2015) to our BowTie and GS datasets as well as using an assembly and alignment free method (Fan *et al.*, 2015) using the total set of reads output by the comparative repeat analysis on the BowTie dataset. Dodsworth et al.’s approach takes into account the assumption that, as repeat abundance changes primarily through random genetic drift (Jurka *et al.*, 2011), they can be used as selection-free characters for phylogenetic reconstruction. On the other hand, AAF methods have been shown to identify potentially useful markers for taxonomic resolution from genome skimming datasets (Bohmann *et al.*, 2020). Both analyses were successful in reconstructing the major relationships between the five *Rhynchospora* species with maximal bootstrap support. Our results not only corroborate that AAF methods can be useful for repetitive sequence data, but also show that it can be employed on the off-target portion of TCS data. These sequence similarity-based approaches (e.g. Vitales et al. 2019) may be more appropriate for groups where no genome skimming data is available for verifying the accuracy of repeat abundance in the target dataset, when abundance-based approaches provide less resolved trees or when genome sizes are unknown.

The phylogenetic relationships retrieved by the repeat abundance and AAF-based phylogeny were not only congruent with a Bayesian analysis of the 256 target regions, but also with recent studies in the genus based on other phylogenetic data (Buddenhagen *et al.*, 2016; Ribeiro *et al.*, 2018). Our results show the potential of repeats from off-target reads to be used as an additional phylogenetically informative dataset, from a completely different part of the genome typically used for phylogenetic studies. Target capture-based sequencing already offers the opportunity to construct robust phylogenies with hundreds of informative markers and also to assemble whole plastomes and other organellar DNAs via the off-target reads to infer phylogenetic relationships, and nuclear-organellar discordance (Dodsworth *et al.*, 2019). Repeat-based phylogenies offer an additional strategy, based on the same TCS datasets, potentially uncovering nuclear intragenomic (in)congruence, while further increasing the usefulness of TCS datasets. The robustness of the repetitive DNA information obtained from our target datasets can prove useful in a variety of phylogenomic approaches, such as similarity-based repeat phylogenies (Vitales *et al.*, 2019) as well as the alignment-free and abundance-based methods presented here. The ability to characterise genome composition and develop cytogenetic markers from TCS-derived repeat data add further value to these datasets, and insight into the genomic processed underpinning evolution in plants.

## ACKNOWLEGMENTS

This study was supported in part by the Coordenacão de Aperfeicoamento de Pessoal de Nivel Superior-Brasil (CAPES, Finance Code 001), CAPES-PRINT project number 88887.363884/2019-00 (LC), and CNPq (Conselho Nacional de Desenvolvimento Científico e Tecnologico; grant number 141037/2018-0 to LC). The authors are also grateful to Dr. Magdalena Vayo (Facultad de Agronomía, Uruguay) for providing comments and suggestions for the manuscript and to Msc. Erton Almeida for the collection of *Rhynchospora cephalotes*.

## AUTHORS CONTRIBUTION

LC and APH conceived the ideas and designed the study with important input from AM, GS, AH and SD; LC, AM, CB, BH, WT and VS collected data; LC, AM and GS performed data analysis; LC wrote the manuscript with comments from all the authors.

## DATA AVAILABILITY

The data that supports the findings of this study is openly available in the NCBI GenBank at [www.ncbi.nlm.nih.gov] under BioProjects PRJEB9643, PRJNA672922 and PRJNA672127.

## Supporting Information

**Table S1 –** Detailed annotation of repetitive elements at lineage level for all datasets on all *Rhynchospora* species.

**Table S2 –** Name and genomic abundance of the satellites found in all datasets of the five *Rhynchospora* species.

**Figure S1 –** Barplots of genomic abundance of unclassified repeats in every dataset of all analysed *Rhynchospora* species.

**Figure S2 –** Dotplot comparison of satellites found in all datasets of all analysed *Rhynchospora* species

