## Supporting Information for "Aiming off the target: studying repetitive DNA using target capture sequencing reads"

**Fig. S1** Barplots of genomic abundance of unclassified repeats in every dataset of *R. cephalotes* (a), *R. exaltata* (b), *R. globosa* (c), *R. pubera* (d) and *R. tenuis* (e).

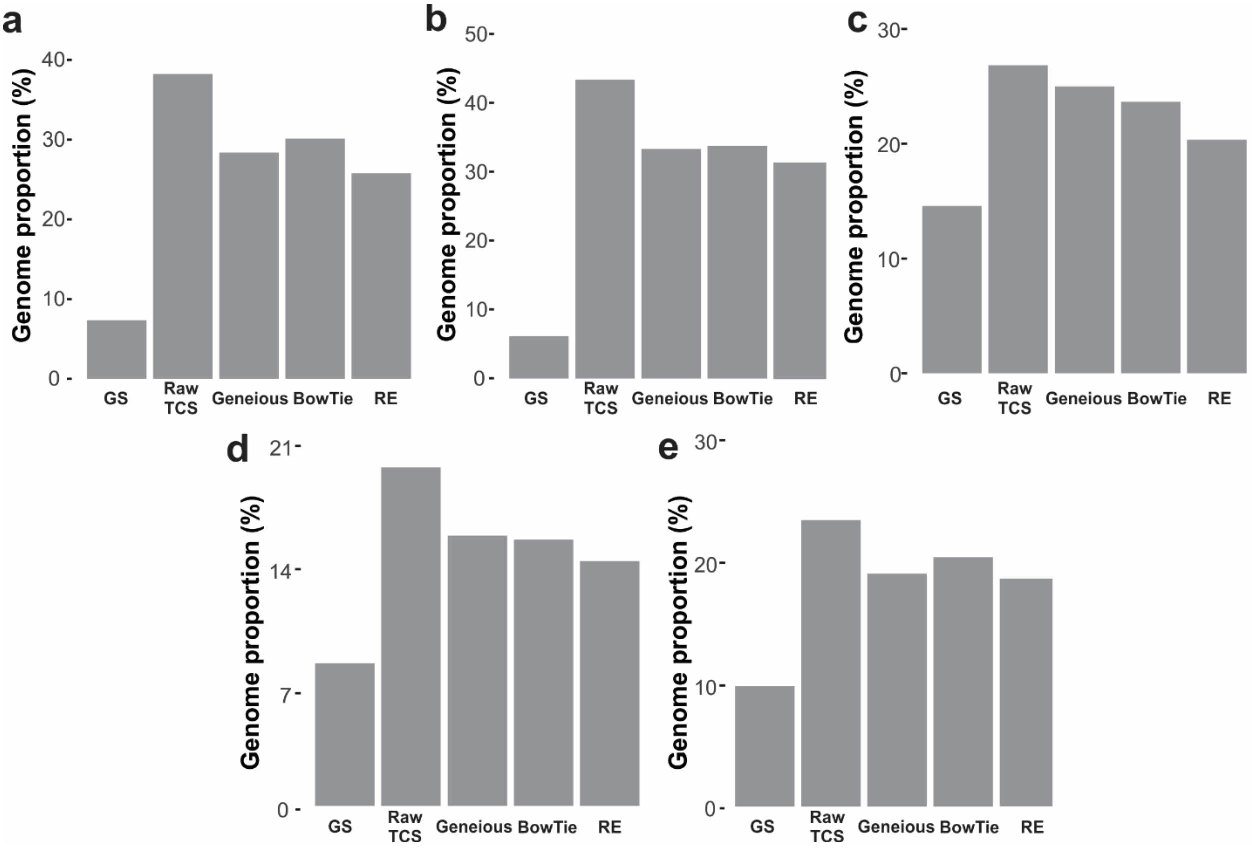


**Fig. S2** Dotplot comparison of all satellites found in the GS (genome skimming, black names), raw TCS (raw target capture, red names), BowTie (blue names), and Geneious (green names) of *R. cephalotes* (A), *R. exaltata* (B), *R. globosa* (C), *R. pubera* (D) and *R. tenuis* (E).

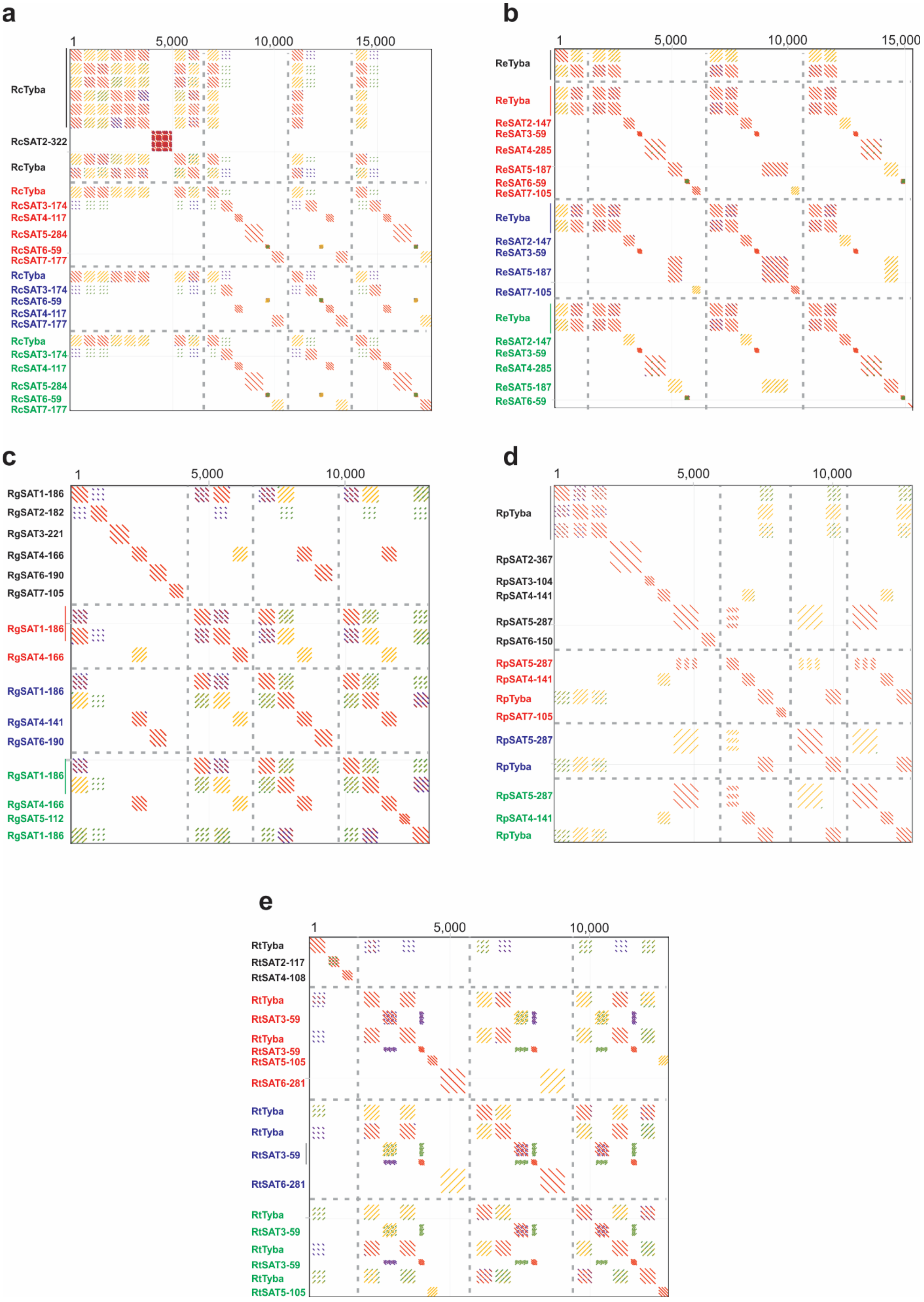
a
1
5,000
10,000
15,000
RcTyba
RcSAT2-322
RcTyba
RcTyba
RcSAT3-174
RcSAT4-117
RcSAT5-284
RcSAT6-59
RcSAT7-177
RcTyba
RcSAT3-174
RcSAT6-59
RcSAT4-117
RcSAT7-177
RcTyba
RcSAT3-174
RcSAT4-117
RcSAT5-284
RcSAT6-59
RcSAT7-177
b
1
5,000
10,000
15,000
ReTyba
ReTyba
ReSAT2-147
ReSAT3-59
ReSAT4-285
ReSAT5-187
ReSAT6-59
ReSAT7-105
ReTyba
ReSAT2-147
ReSAT3-59
ReSAT5-187
ReSAT7-105
ReTyba
ReSAT2-147
ReSAT3-59
ReSAT4-285
ReSAT5-187
ReSAT6-59
c
1
5,000
10,000
RgSAT1-186
RgSAT2-182
RgSAT3-221
RgSAT4-166
RgSAT6-190
RgSAT7-105
RgSAT1-186
RgSAT4-166
RgSAT1-186
RgSAT4-141
RgSAT6-190
RgSAT1-186
RgSAT4-166
RgSAT5-112
RgSAT1-186
d
1
5,000
10,000
RpTyba
RpSAT2-367
RpSAT3-104
RpSAT4-141
RpSAT5-287
RpSAT6-150
RpSAT5-287
RpSAT4-141
RpTyba
RpSAT7-105
RpSAT5-287
RpTyba
RpSAT5-287
RpSAT4-141
RpTyba
e
1
5,000
10,000
RtTyba
RtSAT2-117
RtSAT4-108
RtTyba
RtSAT3-59
RtTyba
RtSAT3-59
RtSAT5-105
RtSAT6-281
RtTyba
RtTyba
RtSAT3-59
RtSAT6-281
RtTyba
RtSAT3-59
RtTyba
RtSAT3-59
RtTyba
RtSAT5-105

**Table S1** Detailed annotation of repetitive elements at lineage level for all datasets on all *Rhynchospora* species.

| ***R. cephalotes*** | | | | | |
|---|---|---|---|---|---|
| Repeat type/Lineage | GS | Raw TCS | Geneious | BowTie | RE |
| UNCLASSIFIELD | 7.392404 | 38.44978 | 28.5435 | 30.28848 | 25.92024287 |
| rDNA 5S | 0.064996 | 0.10635 | 0.147251 | 0.12636 | 0.128520938 |
| rDNA 35S | 0.401899 | 0.210484 | 0.325503 | 0.301439 | 0.254776283 |
| SATELLITE DNA | 3.761397 | 2.844521 | 2.575951 | 2.520998 | 3.437111245 |
| Unclassified LTR | 0.656913 | 0.505674 | 0.769668 | 0.369095 | 0.611092346 |
| Class I/LTR/Ty1/copia/ALE | 0.306891 | 0.170262 | 0.186174 | 0.215879 | 0.205757079 |
| Class I/LTR/Ty1/copia/ANGELA | 0.115084 | 0 | 0.047534 | 0.054228 | 0 |
| Class I/LTR/Ty1/copia/TORK | 0.019876 | 0.0196 | 0.031172 | 0.018248 | 0.02368575 |
| Class I/LTR/Ty1/copia/BIANCA | 0.04174 | 0 | 0 | 0 | 0 |
| Class I/LTR/Ty1/copia/IKEROS | 0.515592 | 0.082319 | 0.103851 | 0.126188 | 0.099480149 |
| Class I/LTR/Ty1/copia/IVANA | 0 | 0 | 0.010506 | 0 | 0 |
| Class I/LTR/Ty1/copia/SIRE | 0 | 0.057095 | 0.066306 | 0.075231 | 0.068997619 |
| Class I/LTR/Ty1/copia/TAR | 0 | 0 | 0.033584 | 0 | 0 |
| LTR-Ty3_gipsy/ATHILA | 0.771202 | 0.243378 | 0.290025 | 0.470492 | 0.294115224 |
| Class I/LTR/Ty3/gypsy/chromovirus/Reina | 0 | 0 | 0.019117 | 0 | 0 |
| Class I/LTR/Ty3/gypsy/non-chromovirus/RETAND | 0 | 0.206224 | 0 | 0 | 0 |
| Class I/LTR/Ty3/gypsy/chromovirus/TEKAY | 0.052871 | 0 | 0 | 0 | 0 |
| Class I/LTR/Ty3/gypsy/non-chromovirus/Tat/Ogre | 0.121444 | 0 | 0.011539 | 0 | 0 |
| Class I/non-LTR/Pararetrovirus | 0.293375 | 0.045165 | 0.078534 | 0.127221 | 0.054580206 |
| Class I/non-LTR/LINE | 0 | 0.057095 | 0 | 0.014633 | 0 |
| Class II/Subclass I/TIR/MuDR_MUTATOR | 0.263759 | 0.124586 | 0.185141 | 0.157347 | 0.150558984 |
| Class II/Subclass I/TIR/EnSpm_CACTA | 0.258591 | 0.104987 | 0.112118 | 0.107251 | 0.126873234 |
| Class II/Subclass I/TIR/hAT | 0.059033 | 0.010396 | 0.027556 | 0 | 0.012563746 |
| Class II/Subclass II/Helitron | 0.034982 | 0 | 0 | 0 | 0 |
| ***R. exaltata*** | | | | | |
| Repeat type/Lineage | GS | Raw TCS | Geneious | BowTie | RE |
| UNCLASSIFIELD | 6.21209 | 43.62766 | 33.52828 | 33.95855 | 31.53302565 |
| rDNA 5S | 0.104813 | 0.099191 | 0.16343 | 0.110793 | 0.120246418 |
| rDNA 35S | 2.136043 | 1.10048 | 1.305278 | 1.042877 | 1.201011929 |
| SATELLITE DNA | 16.10136 | 0.907774 | 3.693127 | 3.390573 | 3.481337117 |
| Unclassified LTR | 0 | 0.5655 | 0.646252 | 0.596279 | 0.646251884 |
| Class I/LTR/Ty1/copia/ALE | 0.022321 | 0.049269 | 0.041825 | 0.018343 | 0.041824841 |

| | | | | | |
|---|---|---|---|---|---|
| Class I/LTR/Ty1/copia/ALESIA | 0 | 0.01417 | 0.015394 | 0.010028 | 0.015393865 |
| Class I/LTR/Ty1/copia/ANGELA | 1.91186 | 0.982322 | 1.421173 | 1.464772 | 1.421173245 |
| Class I/LTR/Ty1/copia/IKEROS | 0.200891 | 0.31 | 0.371486 | 0.650331 | 0.371485915 |
| Class I/LTR/Ty1/copia/IVANA | 0 | 0.051013 | 0.058961 | 0.096363 | 0.058961408 |
| Class I/LTR/Ty1/copia/SIRE | 0 | 0.028122 | 0.03224 | 0 | 0.032239982 |
| LTR-Ty3_gipsy/ATHILA | 0.036879 | 0.561358 | 0.618078 | 0.643727 | 0.618078207 |
| Class I/LTR/Ty3/gypsy/chromovirus/CRM | 0 | 0.01417 | 0.015394 | 0 | 0.015393865 |
| Class I/LTR/Ty3/gypsy/chromovirus/Reina | 0 | 0.245036 | 0.134478 | 0.04598 | 0.134478482 |
| Class I/LTR/Ty3/gypsy/non-chromovirus/RETAND | 0.399841 | 0.770205 | 0.885292 | 0.770662 | 0.885292469 |
| Class I/LTR/Ty3/gypsy/chromovirus/TEKAY | 0 | 0.133418 | 0.338665 | 0.170226 | 0.338665032 |
| Class I/non-LTR/Pararetrovirus | 0 | 0 | 0.014523 | 0 | 0.014522514 |
| Class I/non-LTR/LINE | 0.067934 | 0.129058 | 0.15452 | 0.09245 | 0.154519552 |
| Class II/Subclass I/TIR/MuDR_MUTATOR | 0.054347 | 0.118594 | 0.358706 | 0.46005 | 0.358706102 |
| Class II/Subclass I/TIR/EnSpm_CACTA | 0 | 0.158052 | 0.147549 | 0.147725 | 0.147548745 |

***R. globosa***

| Repeat Type/Lineage | GS | Raw TCS | Geneious | BowTie | RE |
|---|---|---|---|---|---|
| UNCLASSIFIELD | 14.69587 | 27.04666 | 25.19573 | 23.84443 | 20.51788651 |
| rDNA 5S | 0.037092 | 0.10617 | 0.102952 | 0.117128 | 0.116447201 |
| rDNA 35S | 0.447608 | 0.76382 | 0.792483 | 0.692716 | 0.837757591 |
| SATELLITE DNA | 4.189369 | 1.631293 | 1.862184 | 1.744358 | 1.789202971 |
| Unclassified LTR | 3.247036 | 0.844328 | 1.07623 | 0.391098 | 0.926058787 |
| Class I/LTR/Ty1/copia/ALE | 0.421543 | 1.296682 | 1.564878 | 1.294947 | 1.422201128 |
| Class I/LTR/Ty1/copia/ALESIA | 0 | 0.013083 | 0.012053 | 0 | 0.014348944 |
| Class I/LTR/Ty1/copia/ANGELA | 0.24661 | 0.220894 | 0.038168 | 0.293575 | 0.242276405 |
| Class I/LTR/Ty1/copia/TORK | 0 | 0.010567 | 0.038168 | 0.025638 | 0.011589532 |
| Class I/LTR/Ty1/copia/IKEROS | 1.041578 | 0.744196 | 0.799012 | 0.774656 | 0.816234175 |
| Class I/LTR/Ty1/copia/IVANA | 0.0406 | 0.166551 | 0.278223 | 0.187506 | 0.182673098 |
| Class I/LTR/Ty1/copia/SIRE | 0.174933 | 0.225926 | 0.232522 | 0.316699 | 0.24779523 |
| LTR-Ty3_gipsy/ATHILA | 7.973735 | 7.36296 | 8.451645 | 8.03913 | 8.0756962 |
| Class I/LTR/Ty3/gypsy/chromovirus/CRM | 0.01203 | 0.031197 | 0.040679 | 0.010054 | 0.034216713 |
| Class I/LTR/Ty3/gypsy/chromovirus/GALADRIEL | 0 | 0.547454 | 0 | 0 | 0.600448129 |
| Class I/LTR/Ty3/gypsy/chromovirus/Reina | 0 | 0.177621 | 0.026115 | 0.025135 | 0.194814512 |
| Class I/LTR/Ty3/gypsy/non-chromovirus/RETAND | 0.011027 | 0.110699 | 0.20239 | 0.269446 | 0.121414144 |
| Class I/LTR/Ty3/gypsy/chromovirus/TEKAY | 4.49312 | 3.576568 | 6.338357 | 5.211459 | 3.922780605 |
| Class I/LTR/Ty3/gypsy/non-chromovirus/Tat/Ogre | 0.021052 | 0.282281 | 0.417836 | 0.672609 | 0.309606066 |
| Class I/LTR/Ty3/gypsy/non-chromovirus/Tat/TatII | 0 | 0.027675 | 0.018582 | 0.017594 | 0.030353536 |
| Class I/non-LTR/Pararetrovirus | 0.162903 | 0.171583 | 0.195861 | 0.183987 | 0.188191923 |
| Class I/non-LTR/LINE | 0.01203 | 0.121768 | 0.128063 | 0.112604 | 0.133555558 |
| Class II/Subclass I/TIR/MuDR_MUTATOR | 1.098719 | 0.899677 | 1.003912 | 0.832969 | 0.986765858 |
| Class II/Subclass I/TIR/EnSpm_CACTA | 0.573419 | 0.360776 | 0.59612 | 0.547437 | 0.395699732 |

| *R. pubera* | | | | | |
|---|---|---|---|---|---|
| Repeat Type/Lineage | GS | Raw TCS | Geneious | BowTie | RE |
| UNCLASSIFIELD | 9.865057 | 23.47654 | 19.08628 | 20.44017 | 18.66150942 |
| rDNA 35S | 0.607182 | 1.820938 | 1.988511 | 2.039564 | 2.818976485 |
| SATELLITE DNA | 3.869134 | 1.037677 | 1.167537 | 0.907774 | 0.326979111 |
| Unclassified LTR | 2.116154 | 0.483412 | 0.66771 | 0.314338 | 0.513091476 |
| Class I/LTR/Ty1/copia/ALE | 0.432153 | 0.732609 | 0.812793 | 0.768564 | 0.777588542 |
| Class I/LTR/Ty1/copia/ALESIA | 0.01456 | 0.121892 | 0.162537 | 0.132047 | 0.129375795 |
| Class I/LTR/Ty1/copia/ANGELA | 4.509153 | 2.865961 | 3.225208 | 3.117624 | 3.041921451 |
| Class I/LTR/Ty1/copia/TORK | 0.117306 | 0.394385 | 0.434813 | 0.417439 | 0.489493988 |
| Class I/LTR/Ty1/copia/BIANCA | 0.159643 | 0.043885 | 0.04527 | 0.05673 | 0.04657939 |
| Class I/LTR/Ty1/copia/IKEROS | 0.374532 | 0.320921 | 0.342963 | 0.366614 | 0.340624615 |
| Class I/LTR/Ty1/copia/IVANA | 0.025919 | 0.17148 | 0.193735 | 0.192649 | 0.182008454 |
| Class I/LTR/Ty1/copia/SIRE | 1.149722 | 2.092657 | 2.311076 | 2.206849 | 2.221139246 |
| LTR-Ty3_gipsy/ATHILA | 0.510735 | 1.897881 | 2.072398 | 2.275486 | 2.014404728 |
| Class I/LTR/Ty3/gypsy/chromovirus/CRM | 0.011772 | 0.259443 | 0.31231 | 0.289651 | 0.27537243 |
| Class I/LTR/Ty3/gypsy/chromovirus/Reina | 0 | 0.170224 | 0.233442 | 0.148795 | 0.180674683 |
| Class I/LTR/Ty3/gypsy/non-chromovirus/RETAND | 0 | 1.193401 | 1.199608 | 1.028688 | 1.266672139 |
| Class I/LTR/Ty3/gypsy/chromovirus/TEKAY | 0 | 0.460503 | 0.502991 | 0.467295 | 0.488775803 |
| Class I/LTR/Ty3/gypsy/chromovirus/Tcn1 | 0 | 0.141708 | 0 | 0.056923 | 0.150408339 |
| Class I/LTR/Ty3/gypsy/non-chromovirus/Phygy | 0 | 0.114352 | 0 | 0 | 0.121373169 |
| Class I/LTR/Ty3/gypsy/non-chromovirus/Tat/Ogre | 0.852533 | 1.995318 | 2.187482 | 2.067154 | 2.117823286 |
| Class I/LTR/Ty3/gypsy/non-chromovirus/Tat/TatII | 0 | 0 | 0 | 0.045597 | 0 |
| Class I/non-LTR/Pararetrovirus | 0.45993 | 0.459923 | 0.552734 | 0.503889 | 0.488160217 |
| Class I/non-LTR/LINE | 0.215095 | 0.510477 | 0.593423 | 0.549098 | 0.541818853 |
| Class II/Subclass I/TIR/MuDR_MUTATOR | 0.882169 | 0 | 0.539098 | 0.557714 | 0.581421595 |
| Class II/Subclass I/TIR/EnSpm_CACTA | 1.009079 | 0.418937 | 0.467102 | 0.447353 | 0.44465876 |
| Class II/Subclass I/TIR/hAT | 0 | 0.015369 | 0 | 0 | 0.016313046 |
| Class II/Subclass II/Helitron | 0 | 0.067471 | 0 | 0 | 0.071613247 |
| ***R. tenuis*** | | | | | |
| Repeat Type/Lineage | GS | Raw TCS | Geneious | BowTie | RE |
| UNCLASSIFIELD | 8.232814 | 19.52665 | 15.58929 | 15.35823 | 14.12850314 |
| rDNA 5S | 0.021192 | 0.077239 | 0.046696 | 0.078066 | 0.082544154 |
| rDNA 35S | 0.383001 | 0.57813 | 0.683254 | 0.692955 | 0.617835516 |
| SATELLITE DNA | 2.606601 | 3.06471 | 3.454221 | 3.186101 | 3.275192575 |
| Unclassified LTR | 0.373874 | 0.539899 | 0.70668 | 0.660298 | 0.539277398 |
| Class I/LTR/Ty1/copia/ALE | 0.133029 | 0.163493 | 0.180223 | 0.174016 | 0.083706747 |
| Class I/LTR/Ty1/copia/ALESIA | 0 | 0.052529 | 0.075431 | 0.073712 | 0.056136668 |
| Class I/LTR/Ty1/copia/ANGELA | 4.372952 | 1.419682 | 1.439753 | 1.530846 | 1.517184796 |
| Class I/LTR/Ty1/copia/TORK | 0 | 0.019582 | 0 | 0 | 0.020926687 |

| | | | | | |
|---|---|---|---|---|---|
| Class I/LTR/Ty1/copia/BIANCA | 0.100081 | 0.010413 | 0.097295 | 0 | 0.011127683 |
| Class I/LTR/Ty1/copia/IKEROS | 0.968948 | 0.101173 | 0.139774 | 0.135139 | 0.108121215 |
| Class I/LTR/Ty1/copia/IVANA | 0.060173 | 0.108943 | 0.165543 | 0.08973 | 0.116425456 |
| Class I/LTR/Ty1/copia/SIRE | | 0.083611 | 0.120721 | 0.125497 | 0.089353631 |
| LTR-Ty3_gipsy/ATHILA | 0.139062 | 0.321546 | 0.346547 | 0.381312 | 0.343629485 |
| Class I/LTR/Ty3/gypsy/chromovirus/Reina | | 0.068847 | 0.083084 | 0.067958 | 0.073575574 |
| Class I/LTR/Ty3/gypsy/non-chromovirus/RETAND | | 0.156033 | 0.168822 | 0.143692 | 0.166749155 |
| Class I/LTR/Ty3/gypsy/chromovirus/TEKAY | 0.040992 | 0.152148 | 0.078086 | 0.064848 | 0.162597035 |
| Class I/LTR/Ty3/gypsy/chromovirus/Tcn1 | 0 | 0 | 0 | 0 | 0.012907379 |
| Class I/LTR/Ty3/gypsy/non-chromovirus/Tat/Ogre | 0.290344 | 0.022224 | 0.081053 | 0.05334 | 0.023750129 |
| Class I/LTR/Ty3/gypsy/non-chromovirus/Tat/TatII | 0 | 0.014764 | 0.01062 | 0 | 0.015778058 |
| Class I/non-LTR/Pararetrovirus | 0 | 0.038231 | 0.03592 | 0.040588 | 0.040856865 |
| Class I/non-LTR/LINE | 0.033412 | 0.081746 | 0.071839 | 0.05163 | 0.087360613 |
| Class II/Subclass I/TIR/MuDR_MUTATOR | 0.170773 | 0.435462 | 0.516931 | 0.422678 | 0.465369655 |
| Class II/Subclass I/TIR/EnSpm_CACTA | 0.072393 | 0.242597 | 0.278924 | 0.288472 | 0.259258398 |
| Class II/Subclass I/TIR/hAT | | 0.046002 | 0.031391 | 0.173239 | 0.049161106 |
| Class II/Subclass I/TIR/PIF_Harbinger | 0 | 0.012277 | 0 | 0 | 0.0131207 |

**Table S2** Name and genomic abundance of the satellites found in all datasets of the five *Rhynchospora* species.

| **Sp.** | **Satellite** | **Genomic abundance (%)** | | | |
|---|---|---|---|---|---|
| | | **GS** | **Raw TCS** | **BowTie** | **Geneious** |
| ***R. cephalotes*** | *Rc*Tyba | 3.66 | 2.04 | 2.38 | 2.37 |
| | *Rc*SAT2-322 | 0.03 | - | - | - |
| | *Rc*SAT3-174 | - | 0.07 | 0.07 | 0.08 |
| | *Rc*SAT4-117 | - | 0.03 | 0.03 | 0.04 |
| | *Rc*SAT5-284 | - | 0.03 | - | 0.04 |
| | *Rc*SAT6-59 | - | 0.02 | 0.03 | 0.03 |
| | *Rc*SAT7-177 | - | 0.01 | 0.01 | 0.01 |
| ***R. exaltata*** | *Re*Tyba | 16.24 | 2.31 | 2.73 | 2.92 |
| | *Re*SAT2-147 | - | 0.24 | 0.37 | 0.29 |
| | *Re*SAT3-59 | - | 0.19 | 0.20 | 0.24 |
| | *Re*SAT4-285 | - | 0.14 | - | 0.19 |
| | *Re*SAT5-187 | - | 0.03 | 0.03 | 0.04 |
| | *Re*SAT6-59 | - | 0.01 | - | 0.01 |
| | *Re*SAT7-105 | - | 0.01 | 0.01 | - |
| ***R. globosa*** | *Rg*SAT1-186[a] | 3.44 | 1.46 | 1.51 | 1.67 |
| | *Rg*SAT2-182 | 0.24 | - | - | - |
| | *Rg*SAT3-221 | 0.14 | - | - | - |
| | *Rg*SAT4-166 | 0.12 | 0.07 | 0.07 | 0.08 |
| | *Rg*SAT5-112 | - | - | - | 0.07 |
| | *Rg*SAT6-190 | 0.05 | - | 0.05 | - |
| | *Rg*SAT7-156 | 0.01 | - | - | - |

| | | | | | |
|---|---|---|---|---|---|
| ***R. pubera*** | *Rp*Tyba[b] | 3.21 | 0.12 | 0.14 | 0.14 |
| | *Rp*SAT2-367 | 0.20 | - | - | - |
| | *Rp*SAT3-104 | 0.11 | - | - | - |
| | *Rp*SAT4-141 | 0.10 | 0.16 | - | 0.19 |
| | *Rp*SAT5-287 | 0.09 | 0.67 | 0.70 | 0.75 |
| | *Rp*SAT6-150 | 0.04 | - | - | - |
| | *Rp*SAT7-105 | - | 0.01 | - | - |
| ***R. tenuis*** | *Rt*Tyba[a] | 2.56 | 2.66 | 2.77 | 2.97 |
| | *Rt*SAT2-117 | 0.03 | - | - | - |
| | *Rt*SAT3-59 | - | 0.38 | 0.40 | 0.44 |
| | *Rt*SAT4-108 | 0.01 | - | - | - |
| | *Rt*SAT5-105 | - | 0.01 | - | 0.01 |
| | *Rt*SAT6-281 | - | 0.01 | 0.01 | - |

[a] – Previously described in Ribeiro et al. (2017)

[b] – Previously described in Marques et al. (2015)
